# Free diffusion of PI(4,5)P_2_ in the plasma membrane in the presence of high density effector protein complexes

**DOI:** 10.1101/2022.01.07.475414

**Authors:** Jonathan Pacheco, Anna C. Cassidy, James P. Zewe, Rachel C. Wills, Gerald R. V. Hammond

## Abstract

The lipid phosphatidyl-D-*myo-*inositol-4,5-*bis*phosphate [PI(4,5)P_2_] is a master regulator of plasma membrane (PM) function. It engages effector proteins that regulate diverse traffic, transport, signaling and cytoskeletal processes that define PM structure and function. How a single class of lipid molecules independently regulate so many parallel processes remains an open question. We tested the hypothesis that spatially segregated pools of PI(4,5)P_2_ are associated with, and thus reserved for regulation of, different functional complexes in the PM. The mobility of PI(4,5)P_2_ in the membrane was measured using lipid biosensors by single particle tracking photoactivation localization microscopy (sptPALM). We found that PI(4,5)P_2_, and several other classes of inner PM lipids, diffuse rapidly at approximately 0.3 µm^2^/s with largely Brownian motion, although they spend approximately a third of their time diffusing much more slowly. Surprisingly, areas of the PM occupied by PI(4,5)P_2_-dependent complexes, such endoplasmic-reticulum:PM contact sites, clathrin-coated structures, and several actin cytoskeletal elements including focal adhesions, did not cause a change in PI(4,5)P_2_ lateral mobility. Only the spectrin and septin cytoskeletons were observed to produce a slowing of PI(4,5)P_2_ diffusion. We conclude that even structures with high densities of PI(4,5)P_2_-engaging effector proteins, such as clathrin coated pits and focal adhesions, do not corral free PI(4,5)P_2_, questioning a role for spatially segregated PI(4,5)P_2_ pools in organizing and regulating parallel PM functions.

## Introduction

The inner leaflet of an animal cell’s plasma membrane (PM) is a bustling hub of transport, signaling and structure. It is primarily here that cells regulate incoming and outgoing vesicular traffic, control selective permeability through channels and transporters, and facilitate ion and lipid exchange with the endoplasmic reticulum by maintaining membrane contact sites. The lipid bilayer maintains structural rigidity by attaching the underlying cortical cytoskeleton, and builds adhesion complexes that enable cells to integrate into tissues. It also assembles numerous signal transduction complexes to relay extrinsic signals and modify cell function to meet organismal needs. Regulation of these diverse processes relies on proteins that are recruited to and/or activated at the PM by a single class of master regulatory molecule: the lipid, PI(4,5)P_2_ (Saarikangas et al., 2010; Schink et al., 2015; Hammond and Hong, 2018; Dickson and Hille, 2019; Hammond and Burke, 2020). Therefore, understanding the spatial distribution of PI(4,5)P_2_ in the PM, and how it couples to these manifold proteins, is essential to understanding PM function at large.

An attractive hypothesis has been that PI(4,5)P_2_ is spatially segregated into pools that couple to specific PM functions; these functions can then be independently regulated through local changes in PI(4,5)P_2_ concentration, either through lipid corralling or local metabolism (Gamper and Shapiro, 2007; Hammond, 2016). Indeed, specific enrichment of the lipid has been observed at sites of regulated exocytosis, caveolae and clusters of actively signaling K-Ras4B (Bogaart et al., 2011; Trexler et al., 2016; Zhou et al., 2015; Fujita et al., 2009; Zhou et al., 2021). However, in these cases it is likely that PI(4,5)P_2_-binding proteins are responsible for enriching the lipid; there is no evidence that a pre-existing local PI(4,5)P_2_ pool recruits the proteins, or that physiological changes in local PI(4,5)P_2_ concentration modulate their function. Synthesis of PI(4,5)P_2_ has been reported to localize specifically at lipid rafts (Johnson et al., 2008; Myeong et al., 2021). Could this underpin spatial control of individual PM functions? It is worth noting that the transport, signaling and cytoskeletal processes regulated by PI(4,5)P_2_ occur in complexes that are hundreds of nanometers to microns in size, and happen over second to minute time scales. Lipid rafts, on the other hand, are nanoscopic and ephemeral structures in living cells, resolving over nanometer and millisecond scales (Levental et al., 2020).

If cells have the ability to form and maintain spatially segregated pools of PI(4,5)P_2_, this must occur in the context of opposing diffusion of this molecule in the fluid environment of the PM. PI(4,5)P_2_ diffusion has been found to be rapid, with a diffusion coefficient of 0.1-1 µm^2^/s in living cells (Mashanov and Molloy, 2007; Yaradanakul and Hilgemann, 2007; Golebiewska et al., 2008; Hammond et al., 2009). However, these measurements have been obtained from studies of bulk diffusion in the PM and may not detect reduced mobility in the macromolecular complexes driving PM function. Indeed, diffusion of extracellular lipids has been observed to be slowed by the cortical cytoskeleton’s attachment to the PM (Fujiwara et al., 2002; Andrade et al., 2015; Fujiwara et al., 2016). We therefore tested the hypothesis that PI(4,5)P_2_ diffusion is reduced at macromolecular complexes that drive specific PM functions. To this end, we employed single particle tracking photoactivation localization microscopy, sptPALM (Manley et al., 2008) , to measure the diffusion of PI(4,5)P_2_ in living cells. We determine diffusion at PI(4,5)P_2_-dependent macromolecular complexes labelled by expression of fluorescently tagged transgenes or incorporation of such tags onto endogenous proteins by gene editing. We report that, for the most part, free PI(4,5)P_2_ diffuses unhindered inside and between such complexes. However, PI(4,5)P_2_ diffusion is substantially reduced in regions of the membrane that are highly enriched with spectrin or septin filaments, which are components of the cortical cytoskeleton that integrate tightly with the membrane.

## Results

### Diffusion of inner leaflet PI(4,5)P_2_ and other lipids measured by sptPALM using genetically encoded lipid biosensors

PI(4,5)P_2_ in the PM exists in a dynamic equilibrium with its effector proteins (**figure 1**). Estimates using fluorescent acyl chain derivatives indicate that two out of three PI(4,5)P_2_ molecules are in complex with such proteins (Golebiewska et al., 2008). In this manuscript, we consider the remaining one third of lipid molecules that are estimated to be free. When new complexes are assembled, or new effector proteins are recruited to these complexes, it is this free lipid that recruits them; this is the lipid pool that must be locally concentrated to modulate an effector complex. We therefore measured the diffusion of these free lipid molecules. To this end, we used genetically encoded lipid biosensors, which interact with the headgroup. The biosensors themselves are in rapid dynamic equilibrium with the lipids and preclude interaction with endogenous effector proteins (**figure 1**). Unlike effector proteins, the biosensors are estimated to sequester a much smaller fraction of PI(4,5)P_2_, likely less than 10% (Wills et al., 2018). Crucially, previous studies have shown that the diffusion coefficient of biosensor-bound PI(4,5)P_2_ is unchanged from the free lipid (Mashanov and Molloy, 2007; Yaradanakul and Hilgemann, 2007; Golebiewska et al., 2008; Hammond et al., 2009).

**Figure 1:**
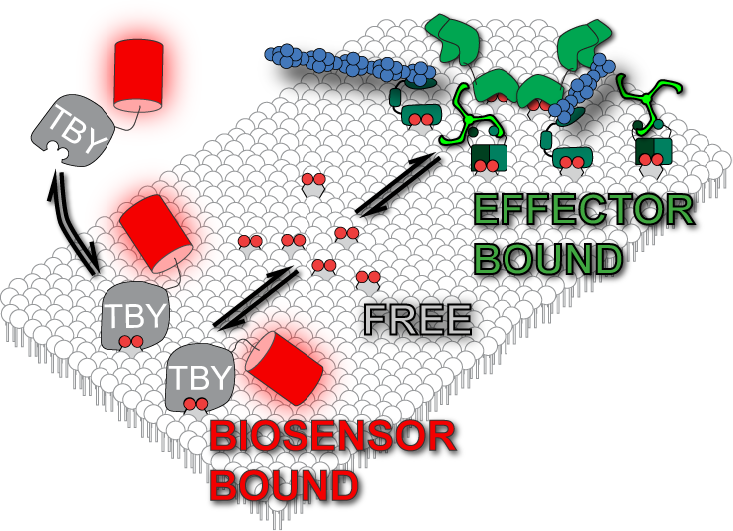
Lipid biosensors and pools of plasma membrane lipid. Functional membrane lipids such as PI(4,5)P_2_ are expected to exist in a dynamic equilibrium between “free lipid” where the headgroup does not engage proteins, and “effector bound” pool where headgroup binds effector proteins. Biosensors like Tubbyc-PAmCherry (TBY) reversibly interact with and thus sample the “free” pool of lipid.

To obtain local diffusion coefficients in intact PM of living HeLa cells, we employed sptPALM (Manley et al., 2008). In this approach, a photoactivatable fluorescent protein is switched on with low intensities of the activating wavelength (in our case, PAmCherry1 with 405 nm light), sufficient to generate sparse and resolvable single fluorescent molecules on the ventral PM, when viewed by total internal reflection fluorescence microcopy (TIRFM; **figure 2A**).

**Figure 2:**
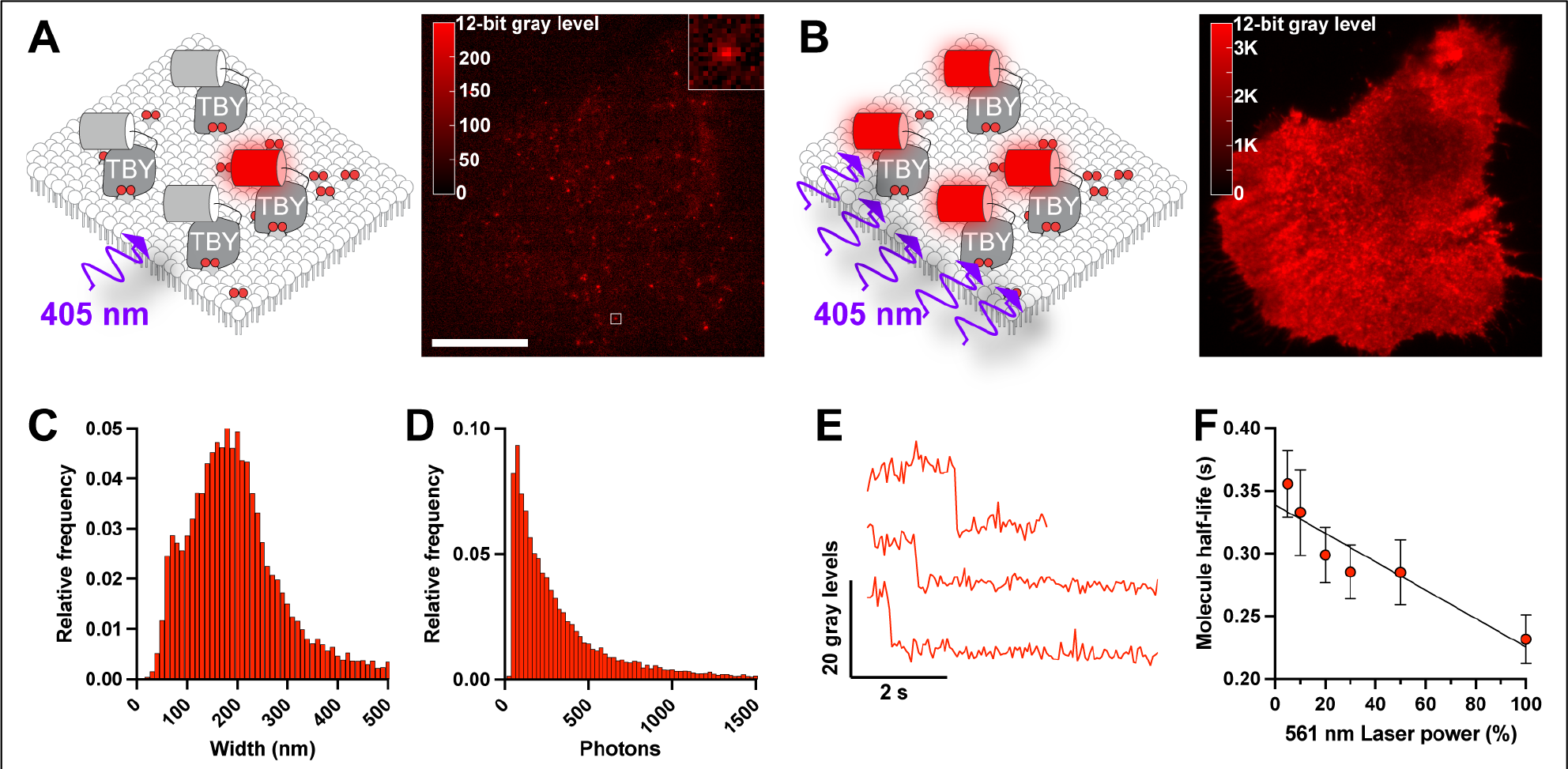
Single molecule detection of lipid biosensors in the PM. (**A**) Single particle tracking PALM using PAmCherry-tagged lipid biosensors. Illumination of a HeLa cell expressing PAmCherry-Tubbyc PI(4,5)P_2_ biosensor with low intensity 405 nm light activates few fluorescent proteins that can be resolved as individual fluorescent spots. Scale bar = 20 μm, inset is 2 μm. (**B**) Activation of PAmCherry with high-intensity 405 nm illumination activates the majority of PAmCherry-Tubbyc, revealing the overall PM distribution in the same HeLa cell as B. (**C**) Fitted diameter and (**D**) photon count of 14,095 individual spots from a representative cell detected with Thunderstorm. (**E**) Example fluorescence intensity profiles of individual spots showing single-step photobleaching. (**F**) Half-life of spot trajectories in time-lapse images measured at varying power modulation of the 561 nm excitation laser, showing half-life decreases proportionally with excitation power; data are grand means ± s.e. from time-lapse recordings of seven cells.

Subsequent activation with high intensities of 405 nm light leads to activation of a large population of molecules, which are no longer resolvable. This leads to the characteristic, uniform sheet of PM fluorescence as seen with non-activatable fluorescent protein conjugates in TIRFM (**figure 2B**). To confirm that single fluorescent puncta were indeed single molecules, we applied the stringent DISH criteria: “1) <u>d</u>iffraction-limited size, 2) intensity of emission appropriate for a single fluorophore, 3) <u>s</u>ingle step photobleaching, and 4) <u>h</u>alf-life of the fluorophore population before photobleaching occurred, [inversely] proportional to laser excitation power” (Mashanov et al., 2004). Indeed, our fluorescent spots had 1) a diffraction limited size, with a mean of 201 nm, consistent with the expected 206 nm size at 596 nm peak emission when imaged through our 1.45 NA objective lens (**figure 2C**); 2) an intensity distribution that is log normal with a mean of 470 photons (**figure 2D**), consistent with prior measurements of PAmCherry1 (Subach et al., 2009); 3) single step photobleaching (**figure 2E**); and 4) a half-life time before photobleaching that was inversely proportional to excitation power (**figure 2F**). Thus, we were able to detect single biosensor:lipid complexes in the PM.

We performed time-lapse imaging of these complexes at approximately 18 frames per second and employed post-hoc tracking analysis using the versatile and accurate single molecule tracking algorithm, TrackMate (Tinevez et al., 2017). This algorithm defines single molecule trajectories, from which we analyzed two properties: the turning angle, θ, between two steps in a trajectory (disregarding direction, giving a range of 0-180°) and the displacement of localizations in the trajectory over increasing time lags (**figure 3A**). We used a variety of lipid biosensors: two for PI(4,5)P_2_, the pleckstrin homology (PH) domain from *PLCD1*, PH-PLC (Várnai and Balla, 1998) and the c-terminal domain from Tubby, Tubbyc; (Quinn et al., 2008); two for the PI(4,5)P_2_ precursor PI4P, which were the PI4P binding domains of *Legionella* effector proteins SidM, P4Mx2; (Hammond et al., 2014) and SidC, P4C (Weber et al., 2014); and one for the abundant inner leaflet phospholipid, phosphatidylserine (PS), namely the C2 domain from lactadherin, Lact-C2 (Yeung et al., 2008). We also employed the myristoylated (C14) and palmitoylated (C16) 11-residue peptide from Lyn kinase, Lyn11 (Teruel et al., 1999). This enabled us to generate a more generalizable measurement of lipid diffusion on the inner leaflet of the PM.

**Figure 3:**
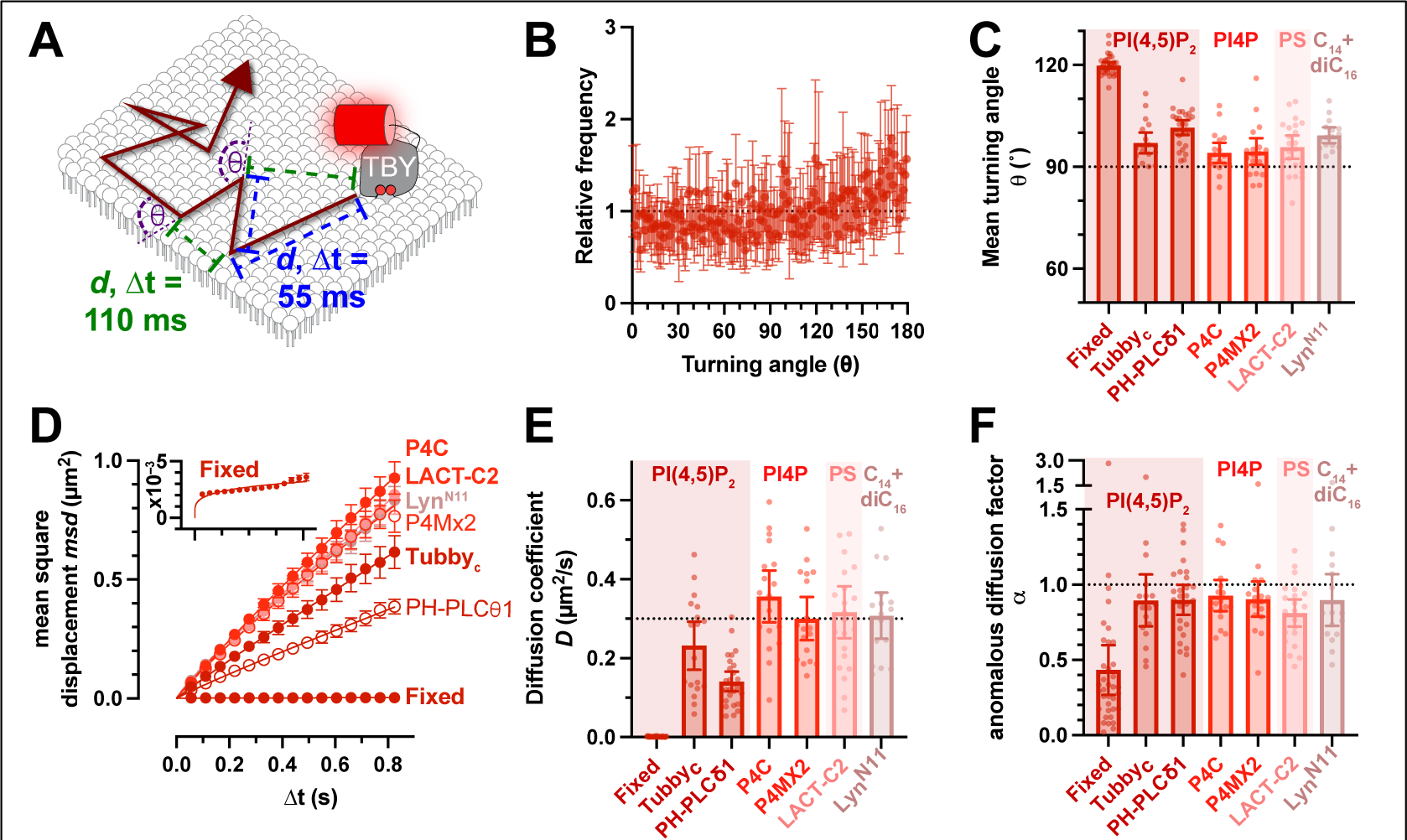
Rapid Brownian diffusion of lipid biosensors in the PM. (**A**) Single particle tracking PALM using PAmCherry-tagged lipid biosensors produces trajectories with measured displacements *d* between localizations separated by various camera exposure-derived time lags, Δ*t*. The turning angle θ between successive displacements can also be measured. (**B**) The distribution of turning angles measured from 18 cells expressing Tubbyc-PAmCherry (means ± 95% C.I.), which is more conveniently represented in (**C**) as the mean turning angle, is close to 90° (no bias in turning angle) for a variety of lipid biosensors. (**D**) These lipid biosensors display a linear increase in mean square displacement over time, indicating Brownian motion. Fit is to *msd* = 4*D*Δtα. (**E**) Mean diffusion coefficients and (**F**) anomalous diffusion factor, α, from individual HeLa cells for the indicated lipid biosensors are shown. The inset for D shows a zoomed axis for the fixed cell data. Data in **C-F** are grand means ± 95% C.I. of trajectories from 18 (Tubbyc, P4C and P4Mx2), 27 (PH-PLCδ1), 22 (Lact-C2), 16 (LynN11) or 35 (Tubbyc-fixed) cells.

For a particle exhibiting true Brownian motion, i.e. unconstrained diffusion, displacement is random, leading to an even distribution of turning angle θ, with a mean of 90° (Burov et al., 2013). As exemplified for the Tubbyc PI(4,5)P_2_ biosensor, we observed θ distributions that were close to random, with a slight tendency towards more obtuse angles (**figure 3B**). All our probes exhibited a small but significant increase in turning angle from the predicted 90° (between 97° and 102°, **figure 3C and table 1**). Therefore, diffusion seemed almost random, but there was a small tendency to reflect back in the direction of motion, perhaps indicating collision with immobile obstacles (Burov et al., 2013). Considering the displacement of the molecules, their mean square displacement increased largely linearly with increasing time lag (**figure 3D**); the slope of this line defines the diffusion coefficient (Einstein, 1905). The non-PI(4,5)P_2_ biosensors exhibited remarkably similar diffusion coefficients of ∼0.3 µm^2^/s (**figure 3E**). For PI(4,5)P_2_, PH- PLCδ1 was about half that at 0.14 µm^2^/s (95% C.I. 0.12-0.17) and Tubbyc somewhat intermediate at 0.23 µm^2^/s (95% C.I. 0.17-0.29). ANOVA revealed a consistent significant difference of PH-PLCδ1 to the other probes, whereas Tubbyc was not consistently significantly different from the other probes (**table 2**). Therefore, we can conclude that the PH-PLCδ1 biosensor diffuses slower than the other lipid probes, but this is not clear for Tubbyc. Notably, our previous work showed that impeded diffusion of the PH-PLCδ1 protein is not a function of its lipid binding properties, and likely represents an additional, undefined interaction(s) that impedes its diffusion (Hammond et al., 2009). On the other hand, our estimates of inner leaflet lipid diffusion here are consistent with prior estimates for PI(4,5)P_2_ that ranged from 0.1-1 µm^2^/s (Mashanov and Molloy, 2007; Yaradanakul and Hilgemann, 2007; Golebiewska et al., 2008; Hammond et al., 2009).

**Table 1:** One sample, 2-tailed t-test of turning angle θ, comparing to a theoretical mean of 90°. Data from Fig, 3C. Significant results are highlighted in bold. n is the number of cells.

| Biosensor: | Fixed | Tubby<sub>c</sub> | PH-PLCδ1 | P4C | P4MX2 | LACT-C2 | Lyn<sup>N11</sup> |
| --- | --- | --- | --- | --- | --- | --- | --- |
| *n* | 35 | 18 | 27 | 18 | 18 | 22 | 16 |
| *t* | 61.85 | 4.814 | 10.88 | 2.925 | 2.416 | 3.524 | 8.506 |
| *P* (2 tailed) | **<0.0001** | **0.0002** | **<0.0001** | **0.0095** | **0.0273** | **0.0020** | **<0.0001** |

**Table 2:** P values from Tukey’s multiple comparison test for biosensor diffusion coefficients presented in Fig. 3E. Significant variation was observed amongst groups by one-way ANOVA (F = 43.14, P <0.0001). Significant results are highlighted in bold.

| Biosensor: | Fixed | Tubby<sub>c</sub> | PH-PLCδ1 | P4C | P4MX2 | LACT-C2 |
| --- | --- | --- | --- | --- | --- | --- |
| Tubby<sub>c</sub> | **<0.0001** |  |  |  |  |  |
| PH-PLCδ1 | **<0.0001** | 0.0558 |  |  |  |  |
| P4C | **<0.0001** | **0.0054** | **<0.0001** |  |  |  |
| P4MX2 | **<0.0001** | 0.3916 | **<0.0001** | 0.6421 |  |  |
| LACT-C2 | **<0.0001** | 0.1221 | **<0.0001** | 0.8735 | 0.9989 |  |
| Lyn<sup>N11</sup> | **<0.0001** | 0.3050 | **<0.0001** | 0.8001 | >0.9999 | >0.9999 |

For freely diffusing particles, the apparent diffusion coefficient *D* is constant at all observed time lags. However, impediments to free diffusion can cause the apparent *D* to decrease at increasing time lags, leading to a downward curve in the mean square displacement vs time plot; this curve is described by the exponent α, where α = 1 describes free diffusion and α < 1 describes impeded or anomalous diffusion (Saxton, 1994). Our range of measurements for α was close to 1 for all probes (**figure 3F**), for the most part averaging ≥ 0.9, which is considered Brownian motion. The one exception was Lact-C2 at α = 0.81 (95% C.I. = 0.72-0.90), which is significantly reduced from 1 (one sample t-test, hypothesized mean = 1, t=3.524, *P* = 0.002).

Nonetheless, deviations from Brownian motion tended to be the exception for these lipid biosensors, indicating largely free Brownian motion in the inner leaflet of the PM.

As a control for the precision of our single particle tracking experiments, we fixed cells expressing Tubbyc with glutaraldehyde to immobilize the biosensor protein. Analysis of time-lapse data from such cells revealed a highly skewed distribution of turning angle θ towards obtuse angles (**figure 3C**), and a very slow diffusion coefficient (figure 3D-E) of 0.0013 µm^2^/s that was highly anomalous (α = 0.43, 95% C.I. = 0.27-0.60; one-sample t-test comparing to hypothesized value of 1; t = 61.85, *P* < 0.0001). This result did not derive from slow diffusion of the fixed probe; rather, it was driven by a mean square displacement that average 0.0027 µm^2^ across all time lags (see **inset** of **figure 3D**). This corresponds to a constant displacement of √0.0027 µm^2^ = 52 nm, representing the precision of our single molecule localization measurements. This places a limit of lateral resolution on our diffusion measurements of ∼100 nm by the Nyquist sampling theorem.

Analyzing diffusion based on whole trajectories gives a mean diffusion coefficient for that particle, but it does not take into account potential changes in *D* as the molecule encounters obstacles that slow its motion. Indeed, visual inspection of our trajectories seems to show different classes: rapid diffusers with large displacements, slow diffusers with short displacements, and a mixture of large and small displacements, which represented the majority (**figure 4A**). We therefore took an alternative approach to define *D* at the population level. We pooled displacements across entire populations from single cells at specific time lags (e.g. 220 ms, as shown in **figure 4B**). Plotting the cumulative distribution allows a fit of the apparent diffusion coefficient (Vrljic et al., 2002). Fitting a single population of diffusers revealed the experimental population had a higher number of shorter displacements and fewer longer displacements than predicted (**figure 4C**, dashed line). Assuming two populations, a fast and a slow, yielded a much tighter fit to the data (**figure 4C**, solid line). All the lipid biosensors exhibited similar distributions with ∼30% of displacements representing slow diffusion and ∼70% being fast, except PH-PLCδ1 which had a 50/50 split. The fast population *D* ranges from 0.3-0.7 µm^2^/s, which is still consistent with the range estimated for PI(4,5)P_2_ diffusion (Mashanov and Molloy, 2007; Yaradanakul and Hilgemann, 2007; Golebiewska et al., 2008; Hammond et al., 2009), whereas the slow population *D* is < 0.05 µm^2^/s (**figure 4D**). Performing this analysis across the first four time-lags allows the time-dependence of *D* to be interrogated. Across all biosensors, the fast population exhibited diffusion coefficients that were time-lag independent, i.e. α = 1 or very close to this value (**figure 4F**). On the other hand, the slow populations of PH- PLC1δ, Lact-C2, and Lyn11 exhibited α ≤ 0.7, which is significantly different from 1, indicating some degree of anomalous, i.e. hindered, diffusion (**table 3**).

**Figure 4:**
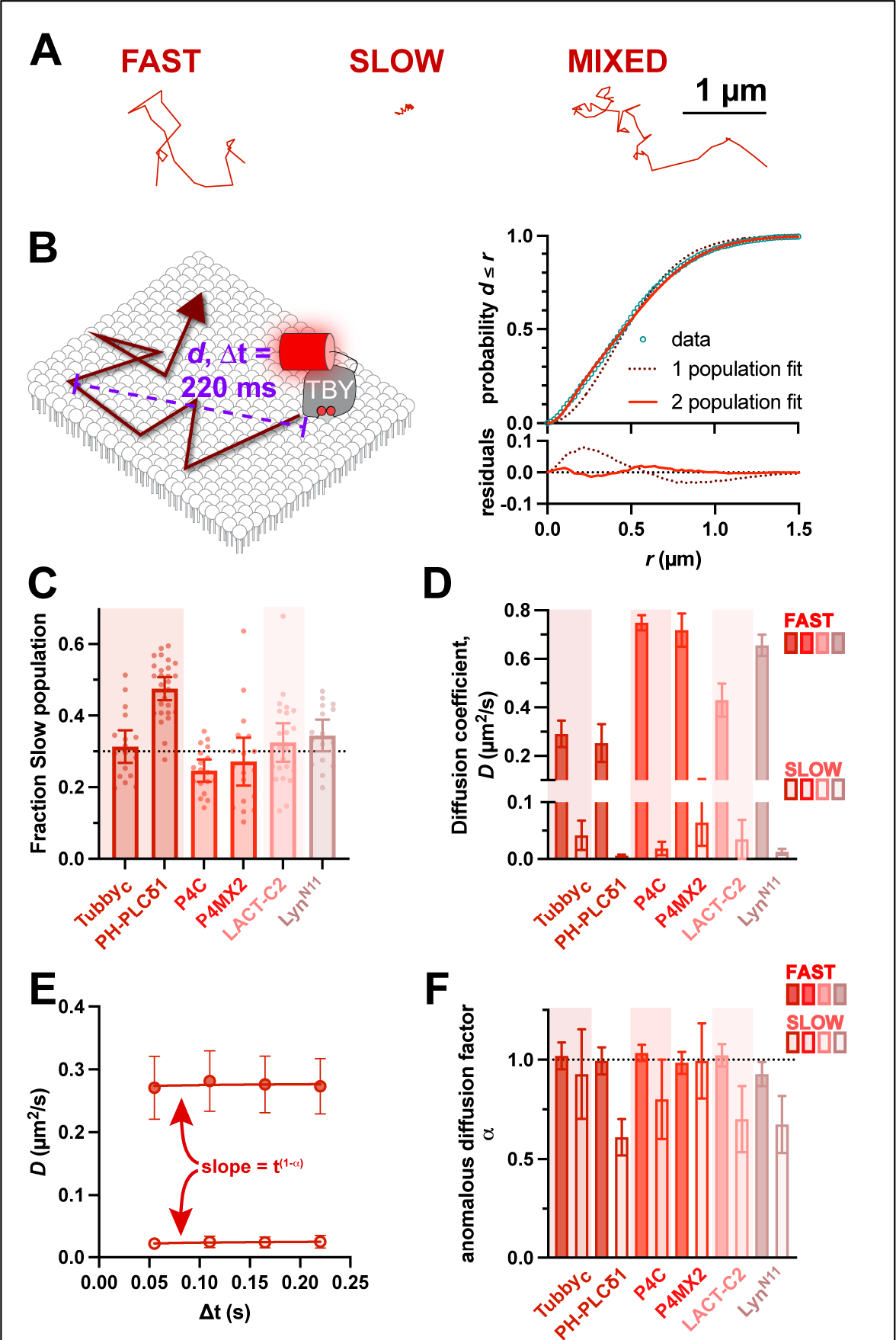
Existence of both fast and slow diffusing lipid molecules in the PM. (**A**) Examples of representative single molecule trajectories, showing either fast moving, slowly moving or mixed trajectories. (**B**) Separating trajectories from all molecules into distinct radial displacements (*d*) at defined Δt (e.g. 220 ms) allows *D* to be estimated from the distribution of *d* values independently of individual (often mixed) trajectories. This distribution is much more tightly estimated by assuming two populations, one fast and one slow. (**C**) Fraction of radial displacements assigned to the slowly diffusing population. (**D**) Mean diffusion coefficients for both fast and slow populations of each lipid biosensor. (**E**) By comparing *D* values from the distribution of radial displacements at different Δt values (i.e. 55, 110, 165 and 220 ms), the dependence of *D* on time interval can be estimated as the slope t1-α, where α = 1 reveals no change and α < 1 indicates decreasing apparent *D* with time. Data are from the Tubbyc biosensor and are grand means ± 95% C.I. (18 cells). (**F**) Anomalous diffusion factor α for both fast (closed) and slow (open symbols) for each biosensor. Only slowly diffusing molecules show evidence of anomalous diffusion. For B, C and E, data are grand means ± 95% C.I. of measurements from the same 18 (Tubbyc, P4C and P4Mx2), 27 (PH-PLCδ1), 22 (Lact-C2) or 16 (LynN11) cells as shown in figure 2.

**Table 3:** One sample, 2-tailed t-test of anomalous diffusion factor α, comparing to a theoretical mean of 1. Data from Fig, 4F. Significant results are highlighted in bold.

| Biosensor: | Tubby<sub>c</sub> |  | PHPLCδ1 |  | P4C |  | P4MX2 |  | LACT-C2 |  | Lyn<sup>N11</sup> |  |
| --- | --- | --- | --- | --- | --- | --- | --- | --- | --- | --- | --- | --- |
|  | Fast | Slow | Fast | Slow | Fast | Slow | Fast | Slow | Fast | Slow | Fast | Slow |
| *n* | 18 |  | 27 |  | 17 |  | 17 |  | 22 |  | 16 |  |
| *t* | 0.5999 | 0.6809 | 0.1799 | 8.787 | 1.798 | 2.119 | 0.5880 | 0.06272 | 0.8254 | 3.756 | 2.546 | 4.831 |
| *P* | 0.5565 | 0.5051 | 0.8586 | **<0.0001** | 0.0911 | 0.0501 | 0.5647 | 0.9508 | 0.4184 | **0.0012** | **0.0224** | **0.0002** |

Therefore, it appears that most free lipids, including PI(4,5)P_2_, exhibit rapid, unhindered Brownian motion on the inner leaflet of the PM. A minor population exhibits slower and potentially hindered diffusion. We next turned our attention to whether this free and hindered diffusion was associated with specific PI(4,5)P_2_-dependent macromolecular complexes in the PM.

### Diffusion of PI(4,5)P_2_ biosensor at PI(4,5)P_2_-dependent effector complexes

For these experiments, we employed the Tubbyc PI(4,5)P_2_ biosensor, because it is ultimately PI(4,5)P_2_ that we are interested in, and Tubbyc behaves most similarly to the other lipid sensors. We performed sptPALM with PAmCherry1-Tubbyc in HeLa cells expressing EGFP- or sfGFP- conjugated markers of specific PI(4,5)P_2_-dependent PM macromolecular complexes. We then segment trajectories based on whether they contact the domains, defined by thresholding of the fluorescence intensity (see materials and methods for details). We then compare diffusion inside these domains to outside at the single cell level.

Although single molecule localization is super-resolution (estimated at ∼100 nm in our experiments), the GFP signal is still subject to the Raleigh limit of lateral resolution when viewed by TIRFM. The threshold-defined domains are therefore convolved with the point-spread function of GFP, appearing slightly larger than they in fact are. For this reason, we interpret our data specifically to interrogate diffusion inside, and in close vicinity of, these structures.

Nonetheless, as will be seen by the numerous examples depicted in the following, trajectories explored the full area of the threshold-defined domains and were not restricted to their periphery. So, although the point spread function “blurring” of the domains’ periphery may skew the data to include some “outside” diffusion behavior as “inside”, changes within the domains will still be detected and all but the subtlest of changes will be measured.

The first structure that we considered were endoplasmic reticulum (ER) – PM contact sites (**figure 5A**). Here, ER membranes are anchored in proximity to the PM to facilitate ion and lipid exchange between the organelles (Wu et al., 2018). Most tethering factors identified to date utilize an interaction with PI(4,5)P_2_ for PM attachment (Giordano et al., 2013; Sohn et al., 2018; Besprozvannaya et al., 2018). We selected one of the most abundant tethering factors, extended synaptotagmin 1 (E-Syt1), tagged with sfGFP on an endogenous allele, allowing us to observe endogenous ER-PM contact sites (Zewe et al., 2018). Endogenous E-Syt1 exhibits a punctate morphology, often strung along tubule-like distributions (**figure 5B**). These structures were frequently crisscrossed by PI(4,5)P_2_:Tubbyc complexes. Surprisingly, we found no changes in diffusion coefficients, α, turning angle, or the fraction of the population exhibiting slow diffusion (**figure 5C**). In short, E-Syt1-defined ER:PM contact sites seemed to offer no impediment to free PI(4,5)P_2_ diffusion.

**Figure 5:**
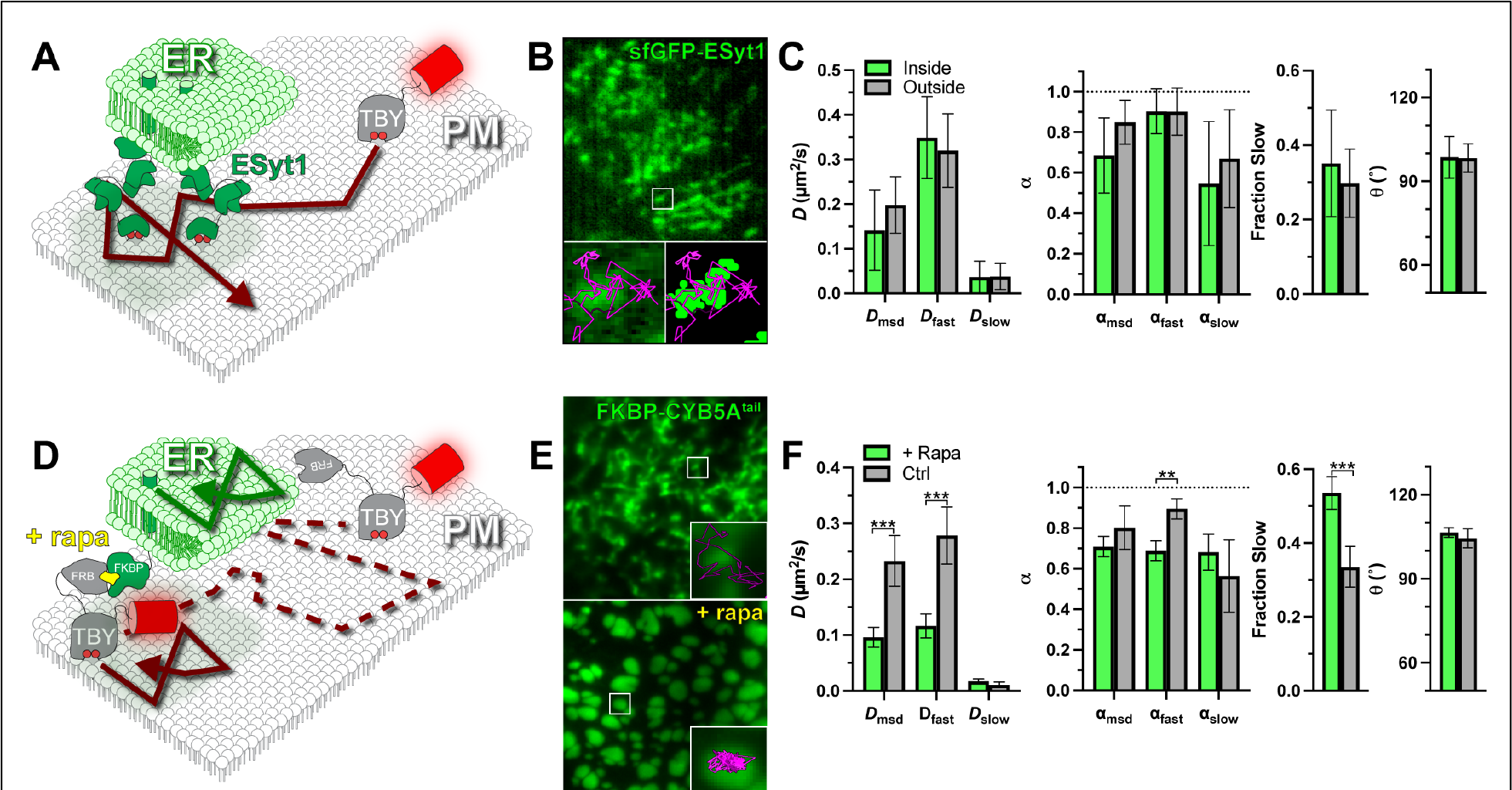
PI(4,5)P_2_ diffusion is uninhibited at ER-PM contact sites. (**A**) Schematic of diffusion in the PM proximal to ER-PM contact sites marked with endogenously tagged sfGFP-ESyt1. (**B**) TIRF image of a small region of PM from an edited sfEGFP-ESyt1 cell; insets show a region of raw and thresholded images overlaid with single molecule trajectories. Inset = 1.5 μm. (**C**) Diffusion coefficients, α (from msd v Δt plots as well as fast and slow populations), the fraction of the population with slow diffusion and mean turning angles are shown for trajectories classified as inside or outside ER-PM contact sites. Data are the grand means ± 95% C.I. of 11 cells. (**D**) FRB-Tubbyc PI(4,5)P_2_ biosensor can be forced into ER-PM contact sites by rapamycin-induced dimerization with ER-resident FKBP-CYB5Atail. (**E**) Images show EGFP-FKBP-CYB5Atail before and after induction of ER-PM contact sites by rapa-induced dimerization with FRB-Tubbyc-PAmCherry. Insets show a region overlaid with single molecule trajectories. Inset = 1.5 μm. (**F**) Diffusion coefficients, α (from msd v Δt plots as well as fast and slow populations), the fraction of the population with slow diffusion and mean turning angles are shown for separate cells treated with or without rapa. Data are grand means ± 95% C.I. of 28 (+rapa) or 19 (control) cells. ** *p* ≤ 0.01, *** *p* ≤ 0.001 from paired t-test with Holm-Šidák correction for multiple comparisons.

To establish that we were able to detect changes in diffusion at ER-PM contact sites, we aimed to trap Tubbyc at ER-PM contact sites using chemically-induced dimerization. To this end, we tagged PAmCherry1-Tubbyc with the FKBP-rapamycin binding (FRB) domain of mTor. We could then recruit this fusion to contact sites using ramapycin to induce FRB dimerization with FK506 binding protein (FKBP) fused to the ER-localized tail of CYB5A (Zewe et al., 2018), as shown in **figure 5D**. When viewed in TIRFM, eGFP-tagged FKBP-CYB5A^tail^ showed characteristic tubular ER morphology before rapamycin addition (pseudocolored green in **figure 5E**), but formed large puncta characteristic of induced ER-PM contact sites upon rapamycin addition (**figure 5E**).

Whereas trajectories of Tubbyc crisscrossed CYB5A-labelled tubules before rapamycin addition, they became trapped within the puncta upon rapamycin addition (**figure 5E**), as expected (**figure 5D**). Notably, we observed substantial changes to diffusion under these conditions.

Upon trapping at contact sites, diffusion halved when assessed either by trajectory mean-square displacements or the fast component of the entire population, and the fast population became slightly, but significantly, more anomalous (**figure 5F**). The fraction of displacements exhibiting slow diffusion also increased from ∼30% to ∼50% (**figure 5F**). On the other hand, diffusion of the slow population was not affected, nor was the distribution of turning angles (**figure 5F**). In a sense, this result was surprising, since trapping inside a domain curtails particle displacements above the domain size, and causes the particles to rebound as they encounter the boundary, increasing the number of obtuse turning angles (Burov et al., 2013). The reason that we did not observe this behavior is likely the large, micron-sized contact sites that were induced (**figure 5E**). Even for the fast population of Tubbyc diffusing at ∼0.1 µm^2^/s imaged across our typical analysis window of 0.22 s, average total displacement will only be 0.17 µm – estimated from √4*D*t/π, where t is the time lag (Teruel and Meyer, 2000). Therefore, the duration of our fluorescence tracking (limited by photobleaching, **figure 2F**) is just below that needed for the edge effects of the large domains to become apparent, explaining why α is only significantly reduced for the fast population (**figure 5F**).

We next turned our attention to clathrin containing structures (CCS). Clathrin-mediated endocytosis is reliant on the PI(4,5)P_2_-dependent recruitment of cargo adapter proteins and fission machinery to build and bud an endocytic vesicle (Mettlen et al., 2018). As a result of this dense, tightly membrane-associated complex, diffusion of unbound lipid has been proposed to be greatly reduced (Schöneberg et al., 2017). We therefore interrogated Tubbyc diffusion in and around CCS by tagging endogenous clathrin light chain with sfGFP (Cho et al.; Leonetti et al., 2016). In these cells, CCS appear as diffraction-limited spots, through which biosensor trajectories passed apparently uninterrupted (**figure 6A, B**). Indeed, we did not measure any reduction in diffusion coefficient or α, nor changes in the turning angle or the fraction of molecules exhibiting slow diffusion (**figure 6C** and **table 4**). Thus, at least for the unbound fraction of PI(4,5)P_2_ at CCS, the assembled clathrin lattice and its network of adaptor proteins do not present a measurable barrier to free diffusion.

**Figure 6:**
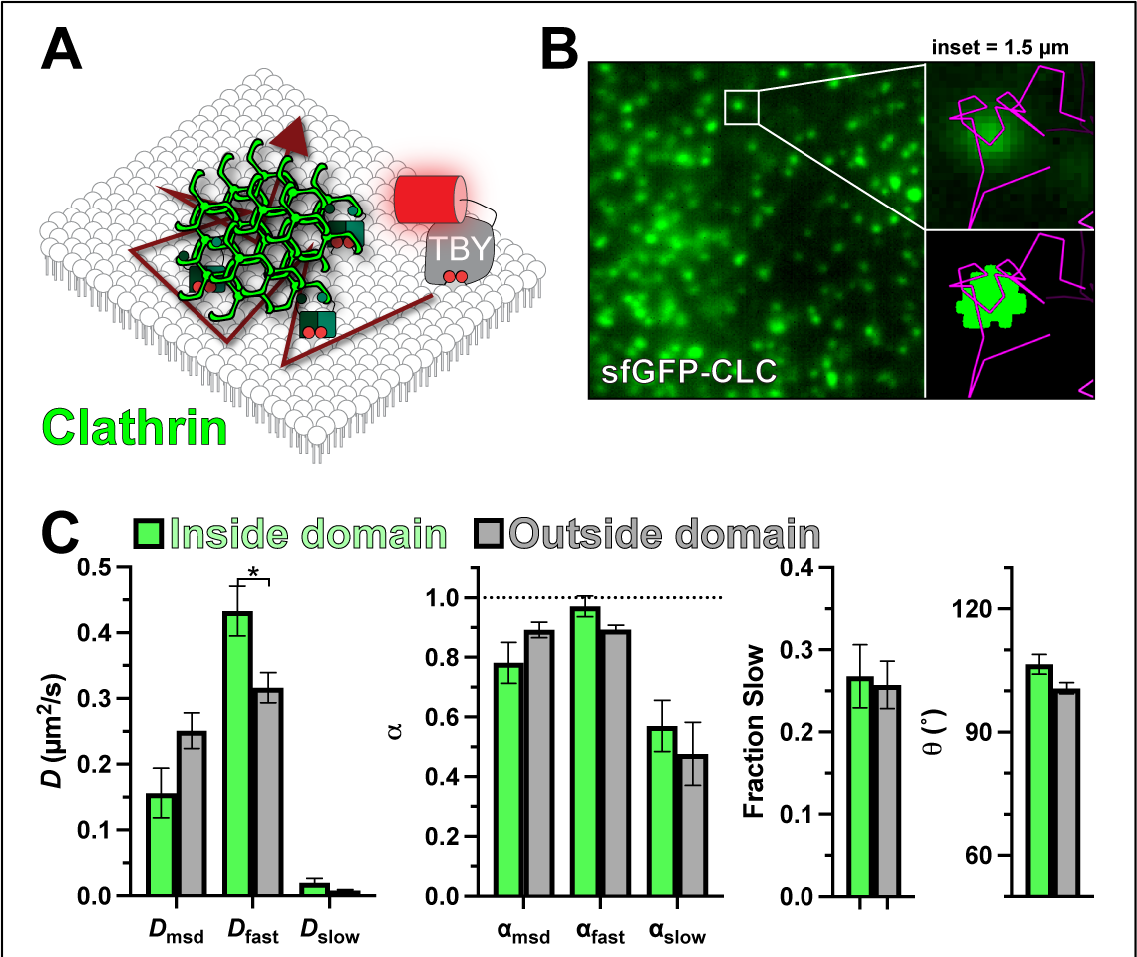
PI(4,5)P_2_ diffusion is uninhibited at clathrin-coated structures. (**A**) Schematic of diffusion in the PM proximal to clathrin coated structures contact sites marked with endogenously tagged sfGFP-Clathrin light chain. (**B**) TIRF image of a small region of PM from an edited sfGFP-CLC cell; insets show a region of raw and thresholded images overlaid with single molecule trajectories. Inset = 1.5 μm. (**C**) Diffusion coefficients, α (from msd v Δt plots as well as fast and slow populations), the fraction of the population with slow diffusion and mean turning angles are shown for trajectories classified as inside or outside ER-PM contact sites. Data are the grand means ± 95% C.I. of 9 cells. * *p* ≤ 0.05 from paired t-test with Holm-Šidák correction for multiple comparisons.

**Table 4:** Results of paired t-tests of the indicated mobility parameters inside and outside domains (or before and after rapamycin addition for FRB-Tubbyc) with the Holm-Šídák correction for multiple comparisons. Data are from Figs. 5- 9. Significant results are highlighted in bold. df is degrees of freedom.

| Domain: |  | ER-PM contact sites | ER-PM contact sites | Clathrin-coated structures | F-actin | Cortical F-actin | Focal Adhesions | Branched F-actin | Spectrin | Septins |
| --- | --- | --- | --- | --- | --- | --- | --- | --- | --- | --- |
| Marker: | | E-Syt1 | FRB-Tubby<sub>c</sub> | CLC | Lifeact | Ezrin | Vinculin | ARPC4 | β-spectrin | Septin-2 |
| Fig: |  | 5A | 5B | 6 | 7 | 7 | 7 | 7 | 8 | 9 |
| $D_{msd}$ | t-ratio | 2.165 | 4.765 | 3.361 | 3.029 | 1.463 | 2.38 | 3.482 | 3.445 | 5.207 |
|  | df | 10 | 18 | 9 | 10 | 13 | 10 | 14 | 15 | 13 |
|  | *P* | 0.3301 | **0.0009** | 0.0571 | 0.0856 | 0.5546 | 0.2700 | **0.0281** | **0.0215** | **0.0014** |
| $D_{fast}$ | t-ratio | 1.512 | 5.161 | 3.721 | 3.656 | 1.775 | 1.231 | 0.5662 | 1.891 | 3.309 |
|  | df | 10 | 18 | 9 | 10 | 13 | 10 | 14 | 15 | 13 |
|  | *P* | 0.5852 | **0.0005** | 0.0375 | **0.0348** | 0.4923 | 0.6773 | 0.8622 | 0.2539 | **0.0224** |
| $D_{slow}$ | t-ratio | 0.103 | 1.215 | 1.665 | 0.02233 | 1.423 | 0.3395 | 1.447 | 1.946 | 0.8402 |
|  | df | 10 | 18 | 9 | 10 | 13 | 10 | 14 | 15 | 13 |
|  | *P* | 0.9995 | 0.5636 | 0.3421 | 0.9826 | 0.5546 | 0.9331 | 0.5254 | 0.2539 | 0.5320 |
| $F_{slow}$ | t-ratio | 1.055 | 5.524 | 0.2427 | 1.196 | 2.087 | 0.1355 | 3.497 | 1.073 | 2.428 |
|  | df | 10 | 18 | 9 | 10 | 13 | 10 | 14 | 15 | 13 |
|  | *P* | 0.7815 | **0.0002** | 0.8137 | 0.7768 | 0.3757 | 0.9331 | **0.0281** | 0.3180 | 0.0886 |
| $\alpha_{msd}$ | t-ratio | 2.349 | 1.371 | 2.011 | 1.529 | 0.3705 | 2.086 | 2.063 | 5.562 | 4.853 |
|  | df | 10 | 18 | 9 | 10 | 13 | 10 | 14 | 15 | 13 |
|  | *P* | 0.2828 | 0.5636 | 0.2895 | 0.6418 | 0.7170 | 0.3256 | 0.3019 | **0.0004** | **0.0022** |
| $\alpha_{fast}$ | t-ratio | 0.08508 | 4.226 | 2.335 | 0.4044 | 1.533 | 1.054 | 0.7198 | 3.102 | 4.251 |
|  | df | 10 | 18 | 9 | 10 | 13 | 10 | 14 | 15 | 13 |
|  | *P* | 0.9995 | **0.0025** | 0.2384 | 0.9715 | 0.5546 | 0.6809 | 0.8622 | **0.0359** | **0.0057** |
| $\alpha_{slow}$ | t-ratio | 1.668 | 0.8014 | 0.7737 | 0.6813 | 1.817 | 1.4 | 2.055 | 1.427 | 1.043 |
|  | df | 10 | 18 | 9 | 10 | 13 | 10 | 14 | 15 | 13 |
|  | *P* | 0.5552 | 0.5636 | 0.7073 | 0.9429 | 0.4923 | 0.6549 | 0.3019 | 0.3180 | 0.5320 |
| $\theta$ | t-ratio | 0.07485 | 1.243 | 2.091 | 0.1832 | 0.8525 | 2.224 | 0.6676 | 4.093 | 4.164 |
|  | df | 10 | 18 | 9 | 10 | 13 | 10 | 14 | 15 | 13 |
|  | *P* | 0.9995 | 0.5636 | 0.2895 | 0.9799 | 0.6512 | 0.3035 | 0.8622 | **0.0067** | **0.0057** |

One PM structure that has been proposed to have profound impacts on lipid diffusion (at least in the outer PM leaflet) is the cortical actin cytoskeleton (Morone et al., 2006; Andrade et al., 2015; Fujiwara et al., 2016). We labelled the endogenous F-actin cytoskeleton with EGFP-Lifeact (Riedl et al., 2008), revealing a dense array of cortical filaments in TIRFM (**figure 7A**). These are thought to be bundled filaments, rather than the meshwork of individual filaments that make up the majority of the cortical or membrane cytoskeleton (Morone et al., 2006). Tubbyc crisscrossed these filaments without noticeable impediment, and we saw no change in any measured parameter to indicate non-Brownian motion in proximity of these filaments (**figure 7A**).

**Figure 7:**
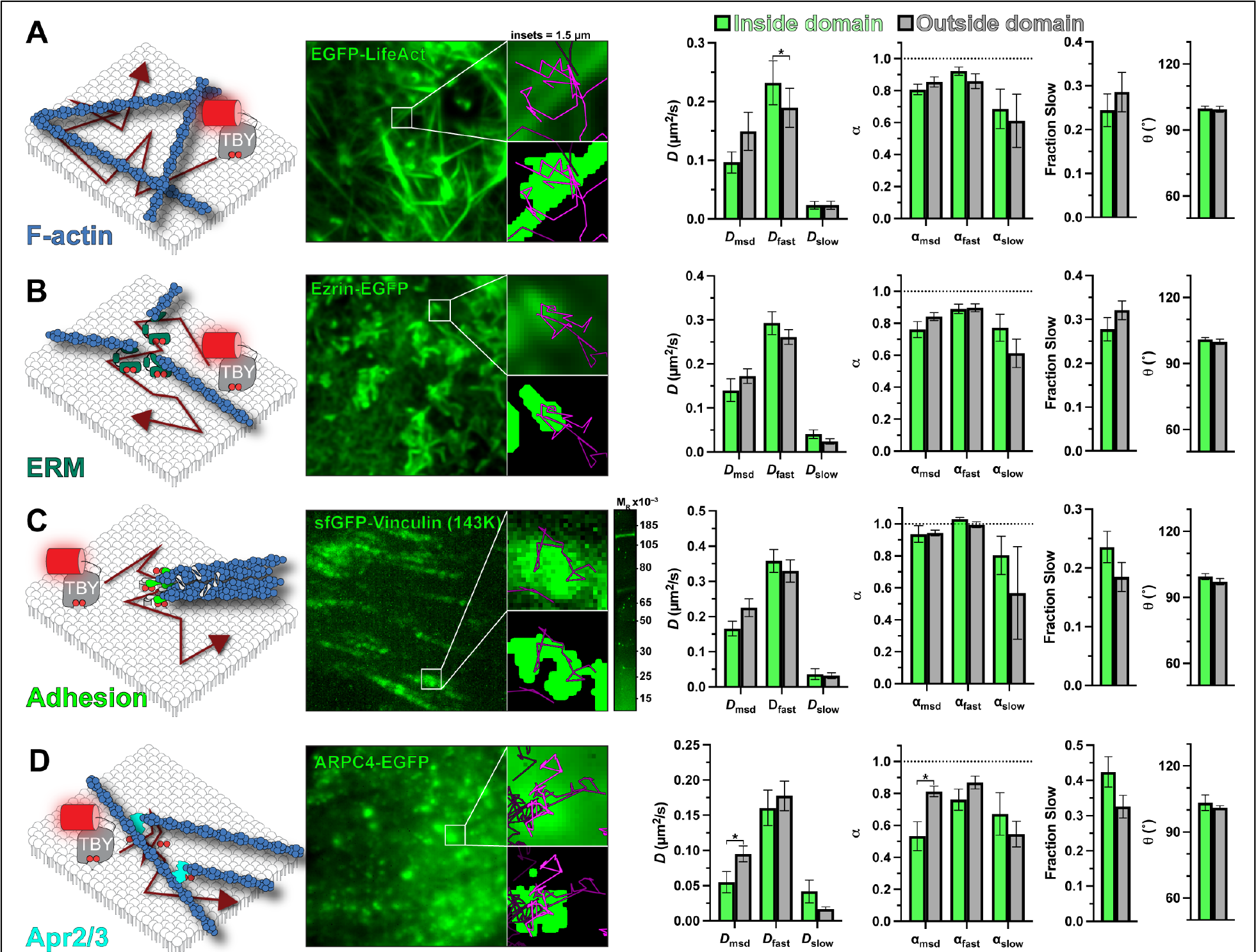
The actin cytoskeleton does not present a substantial impediment to PI(4,5)P_2_ diffusion. Tubbyc- PAmCherry single molecule diffusion was measured inside and outside of the indicated green fluorescent proteinlabeled structure: (**A**) The F-actin cytoskeleton was labelled with LifeAct-GFP. (**B**) Cortical F-actin. Membrane attachment was marked with Ezrin-EGFP expression. (**C**) Focal adhesions were labelled by gene editing the *VCL* locus to express sfGFP-vinculin. (**D**) The ArP_2_/3 complex was labelled by expression of ARPC4-EGFP. In all cases, the images show the GFP-labelled domain, with the 1.5 μm insets showing an isolated domain as the raw image or thresholded image overlaid with molecule trajectories passing through it during the experiment. Data at right show diffusion coefficients and anomalous diffusion (α) from both mean square displacement analysis of trajectories and analysis of individual displacements. The fraction “fast” and “slow” displacements and mean turning angle of trajectories is shown. In all cases, data are the grand means ± s.e. of 11 (A, C), 14 (B) or 15 (D) cells. The gel image in C shows in-gel fluorescence of sfGFP from lysates of VCL-sfGFP cells, with a single band consistent with the expected Mr of 143,000 for the fusion protein. * *p* ≤ 0.05 from paired t-test with Holm-Šidák correction for multiple comparisons.

There is no reason to suspect a specific interaction of PI(4,5)P_2_ with bundled actin filaments. On the other hand, the lipid is intimately involved in activating proteins that attach the F-actin cytoskeleton to the membrane (Saarikangas et al., 2010), so we decided to measure PI(4,5)P_2_ diffusion in proximity to such complexes. We selected three candidates: firstly, we considered ERM-proteins, which anchor actin filaments to the membrane in a PI(4,5)P_2_-dependent manner (Algrain et al., 1993; Senju et al., 2017); these were labelled by expressing Ezrin-EGFP (**figure 7B**). Secondly, we studied focal adhesions, whose assembly is thought to be driven by local PI(4,5)P_2_ synthesis (Legate et al., 2011), activating proteins such as vinculin (Chinthalapudi et al., 2014); this was labelled by integrating a split sfGFP tag at an endogenous *VCL* allele, yielding the expected 143K sfGFP-tagged vinculin protein (**figure 7C**). Thirdly, we considered the branched F-actin nucleating complex ArP_2_/3, which relies on PI(4,5)P_2_ for PM recruitment (Zoncu et al., 2007); we imaged ArP_2_/3 by expressing a tagged component, ARPC4-EGFP (**figure 7D**), which exhibits a punctate PM distribution when imaged by TIRFM (Zoncu et al., 2007), reminiscent of endogenously tagged ARPC3 and 4 in HEK293 cells (Cho et al.). To our surprise, we detected no change in Tubbyc diffusion in and around ERM proteins or focal adhesions (**figure 7B, C** and **table 4**). For ArP_2_/3, we measured a significant decrease in *D* and α measured by mean square displacement (**figure 7D** and **table 4**). This was not reflected by changes in *D* or α for the fast or slow components of diffusion, although there was a roughly 10% increase in the fraction of Tubbyc molecules exhibiting slow diffusion (**figure 7D** and **table 4**), explaining the mean-square displacement result. Collectively, these data do not reveal a substantial impact of the F-actin cytoskeleton on PI(4,5)P_2_ diffusion at the temporal and spatial scales investigated herein – despite a small impediment evident at sites of branched F-actin nucleation by ArP_2_/3.

We next turned our attention to the non-actin components of the cortical cytoskeleton. Spectrins assemble into membrane-proximal filaments that integrate with the F-actin cortex (Bennett and Lorenzo, 2016). Unlike F-actin, these filaments are directly anchored to the PM in a PI(4,5)P_2_- dependent manner (Wang and Shaw, 1995), forming a potential barrier to diffusion (**figure 8A**). We labelled the spectrin cytoskeleton by expressing EGFP-β-spectrin, yielding the expected cortical distribution in HeLa cells when viewed by confocal microscope (**figure 8B**). In TIRFM, a largely amorphous but patchy distribution is observed (**figure 8C**), similar to previous observations in fibroblasts (Ghisleni et al., 2020). Thresholding of this signal therefore identified regions of the PM where spectrin density was highest, with more restricted trajectories (**figure 8C**). Indeed, both the diffusion coefficient and α measured from mean square displacement were reduced, which was also evident specifically in the fast population of molecules (**figure 8D** and **table 4**). The mean turning angle θ was also significantly increased in these regions (**figure 8D** and **table 4**). These observations are consistent with spectrins acting as a diffusion barriers, as high densities of filaments would be expected to reduce the number of long-distance paths to displacement (reducing *D* and α) and cause rebound of PI(4,5)P_2_:Tubbyc complexes that strike them (increasing θ).

**Figure 8:**
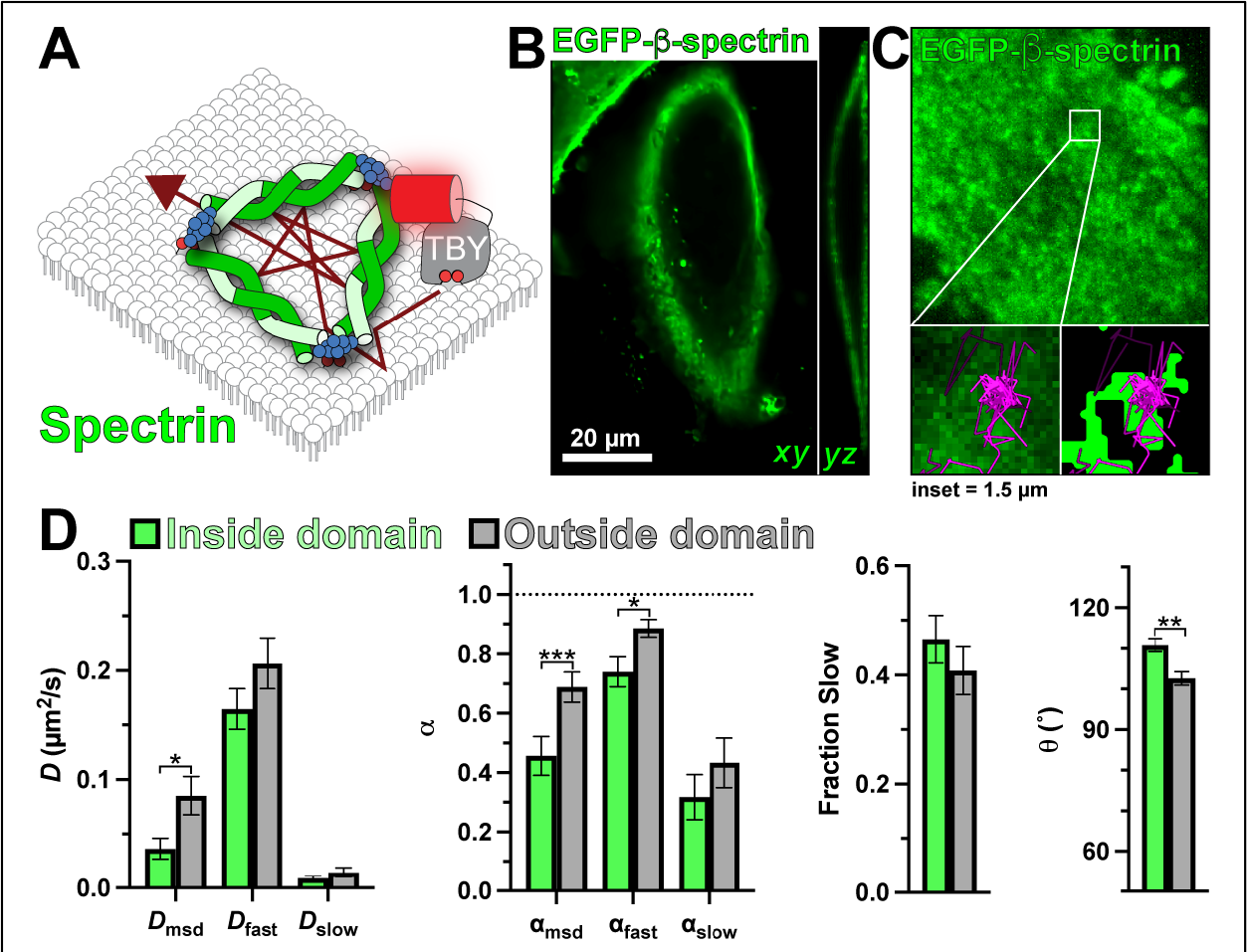
PI(4,5)P_2_ diffusion is impaired by the spectrin cytoskeleton. (**A**) Schematic of diffusion in the PM proximal to spectrin marked with expressed EGFP-β-spectrin. (**B**) Transverse and longitudinal confocal sections and (**C**) TIRF image of a small PM region of HeLa cells expressing EGFP-β-spectrin; insets show a region of raw and thresholded images overlaid with single molecule trajectories. Inset = 1.5 μm. (**D**) Diffusion coefficients, α (from msd v Δt plots as well as fast and slow populations), fraction of the population with slow diffusion and mean turning angles are shown for trajectories classified as inside or outside dense regions of spectrin labelling. Data are the grand means ± s.e. of 15 cells. * *p* ≤ 0.05, ** *p* ≤ 0.01, *** *p* ≤ 0.001 from paired t-test with Holm-Šidák correction for multiple comparisons.

Finally, we interrogated the last component of the cortical cytoskeleton: septins (**figure 9A**). Septin filaments serve as a scaffold for the F-actin and microtubule cytoskeletons at the PM (Spiliotis and Nakos, 2021), and like spectrins, are directly associated with the membrane via an interaction with PI(4,5)P_2_ (Zhang et al., 1999; Tanaka-Takiguchi et al., 2009; Bertin et al., 2010). We edited a split sfGFP tag into the *SEPT2* allele, yielding the expected 63K septin2-sfGFP complex viewed by in-gel fluorescence (**figure 9B**). By TIRFM, these edited HeLa cells exhibited localized filamentous patterns on the ventral PM (**figure 9B**), consistent with a similarly edited cell line from another group (Banko et al., 2019). Strikingly, we observed that although Tubbyc trajectories crossed into, out of, and through such filaments (e.g. **figure 9B**), their mobility was greatly altered; overall diffusion by mean square displacement, and the diffusion of the fast population, were more than halved (**figure 9C**). Diffusion was also significantly more anomalous (α reduced), and the mean turning angle was significantly increased (**figure 9C** and **table 4**).

**Figure 9:**
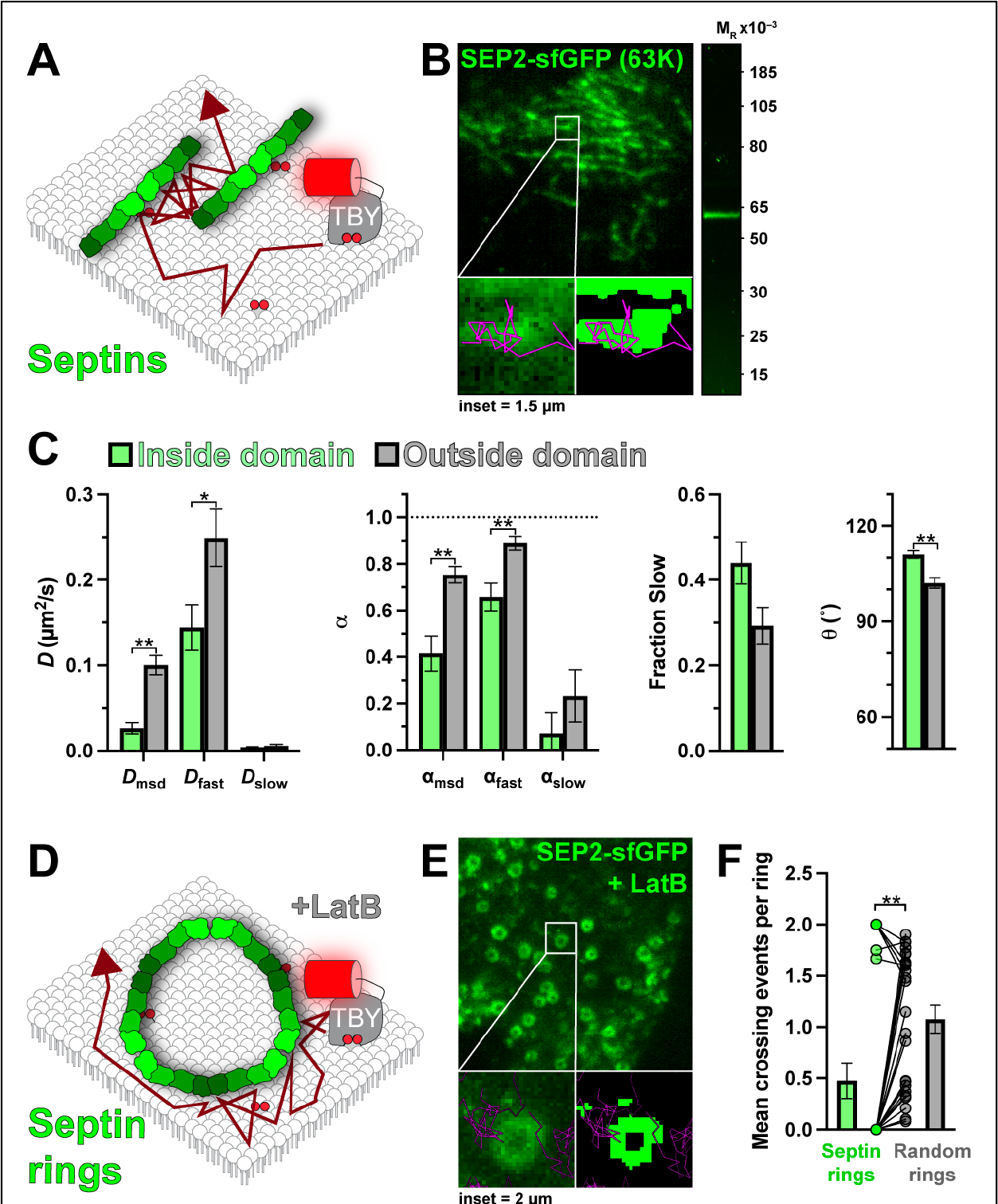
Septins are a barrier to PI(4,5)P_2_ diffusion. (**A**) Schematic of diffusion in the PM proximal to septins marked with endogenous septin2 tagged with sfGFP. (**B**) TIRF image of a small PM region of HeLa septin2-sfGFP cell; insets show a region of raw and thresholded images overlaid with single molecule trajectories. Inset = 1.5 μm. The gel image at right shows in-gel fluorescence of sfGFP from lysates of septin2-sfGFP cells, with a single band consistent with the expected Mr of 63,000 for the fusion protein. (**C**) Diffusion coefficients, α (from msd v Δt plots as well as fast and slow populations), fraction of the population with slow diffusion and mean turning angles are shown for trajectories classified as inside or outside septin filaments. Data are the grand means ± s.e. of 14 cells. * *p* ≤ 0.05, ** *p* ≤ 0.01 from paired t-test with Holm-Šidák correction for multiple comparisons. (**D**) Schematic of diffusion around septin rings induced by treatment of cells with latrunculin B. (**E**) TIRF image of a small PM region of latrunculin Btreated HeLa septin2-sfGFP cell; insets show a region of raw and thresholded images overlaid with single molecule trajectories. Inset = 2 μm. (**F**) Mean number of crossing events into or out of the lumen of septin rings observed in 24 cells is compared to a randomized rings (generated by randomizing the masked rings) shows far fewer crossing events at septin rings. Bars show mean ± s.e., points show pair-wise comparison of individual cells’ septin and randomized rings. ** *p* ≤ 0.01 from Wilcoxon matched-pairs signed rank test.

The simplest explanation of these data is that, like for spectrin, individual septin filaments present a physical barrier to diffusion. However, since the filaments imaged by TIRFM likely represent bundles of many septin filaments, with gaps that cannot be resolved, we sought a more direct test of this hypothesis: we utilized the observation that disruption of the F-actin cytoskeleton causes the septin cytoskeleton to collapse into micron-scale, continuous rings (Xie et al., 1999; Kinoshita et al., 2002). These rings were easily resolved in HeLa cells treated with latrunculin B (**figure 9E**). Strikingly, trajectories tended to explore the periphery of such rings but very rarely crossed into their center. Quantifying the frequency of such entry into the ring lumens revealed such crossing events happened far less frequently than occurred in the same sized, random areas of membrane (**figure 9F**). Therefore, septin2-containing filaments present a bone-fide barrier to diffusion of lipids on the inner leaflet of the PM, as previously suggested (Golebiewska et al., 2011).

## Discussion

Here, we determined the mobility of lipid molecules on the inner leaflet of the PM with ∼100 nm and ∼100 ms resolution, and we interrogated how this mobility changes in proximity to macromolecular complexes regulating diverse PM functions. Broadly speaking, we find diffusion coefficients of approximately 0.3 µm^2^/s (**figures 2-3**), agreeing well with prior estimates of PI(4,5)P_2_ diffusion (Mashanov and Molloy, 2007; Yaradanakul and Hilgemann, 2007; Golebiewska et al., 2008; Hammond et al., 2009), as well as with that of phosphatidylethanolamine in the outer PM leaflet at this level of spatiotemporal resolution (Fujiwara et al., 2002; Andrade et al., 2015). Surprisingly, we found that most PI(4,5)P_2_- regulated macromolecular complexes we investigated did not impact the diffusion of PI(4,5)P_2_ (**figures 5-9**).

One finding that we do not have a completely satisfactory explanation for is the existence of two apparent modes of diffusion in HeLa cells: one rapid and largely Brownian, and another slower and more anomalous (**figure 4**). Notably, these two modes are frequently observed in the same trajectory (**figure 4A**). This observation, when coupled with a slight tendency towards more obtuse turning angles (**figure 3C**), suggests that the inner leaflet of the PM is largely supportive of Brownian motion, but there is a tendency to produce transient, local confinement. Our data do not allow us to unambiguously assign the molecular cause of this confinement. However, we speculate that high local densities of immobile membrane anchored proteins, perhaps associated with the cortical cytoskeleton, may transiently trap the lipids long enough to be detected in our experiments. In HeLa cells, the membrane-associated cortical cytoskeleton constrains lipids in 68 nm corrals (Fujiwara et al., 2016). At the spatiotemporal resolution of our experiments, diffusion coefficients therefore represents the rate of diffusion between these corrals, rather than the more rapid diffusion within them. However, given the variable sizes of such corrals and a stochastic process of escape from them, perhaps our slower population represents lipid molecules trapped within corrals for long enough that the restriction becomes apparent. It is worth mentioning that cortical actin cytoskeletal barriers to lipid diffusion have been shown to be critically dependent on the ArP_2_/3 complex (Andrade et al., 2015), which is the only complex where we observed a significant (albeit small) increase in the fraction of displacements exhibiting the slower mode of diffusion (**table 4** and **figure 4D**).

Whatever the cause of the slower mode of diffusion, it does not seem to be associated with any particular functional complexes probed in this study, since the fraction of displacements assigned to this slower mode was not changed substantially in proximity to any of them, except ArP_2_/3 as discussed above (**figures 5-9, table 4**). This result surprised us, since membrane attached complexes such as clathrin-coated pits and focal adhesions are envisioned to consist of dense arrays of inter-linked, membrane attached proteins (Schöneberg et al., 2017; Kanchanawong et al., 2010). Even for lipid molecules not directly interacting with these proteins, the dense array of anchored proteins would be expected to block long-distance displacements across the complex – i.e., cause percolation of the diffusing lipid. The data presented herein suggest that in fact, the density of membrane-attached proteins is below the percolation threshold for the lipids. Modeling studies have suggested that even with roughly one third of the membrane area occupied by such immobile protein obstacles, impeded diffusion would not be apparent at our spatiotemporal resolution (Saxton, 1994). The implication is that, even though PI(4,5)P_2_ itself can be an anchoring component of the membrane proteins, any unbound PI(4,5)P_2_ can rapidly and freely diffuse away from the complex.

One set of membrane-anchored complexes where we did observe consistent reductions in diffusion were at the spectrin and septin cytoskeleton (**figures 8 & 9**). We could not resolve individual filaments in our TIRFM imaging, so the threshold-defined domains likely represent dense arrays of such filaments. If the filaments are impermeable to lipid diffusion across them, then the convoluted paths between them would explain the slowed and more anomalous diffusion. Direct support for this comes from the induction of resolvable septin rings (**figure 9D**), which lipid trajectories were very rarely able to cross. It therefore seems that septin and spectrin filaments, anchored tightly to the membrane surface by PI(4,5)P_2_ molecules (Wang and Shaw, 1995; Zhang et al., 1999; Tanaka-Takiguchi et al., 2009; Bertin et al., 2010), are indeed true barriers to lipid diffusion, including that of free PI(4,5)P_2_. In support of this, Langevin simulations have revealed such diffusion barriers for septins (Lee et al., 2014).

The exception presented by spectrin and septin filaments aside, our main conclusion is that the endocytic, cytoskeletal, and organelle tethering complexes that we resolved on the inner leaflet of the plasma membrane do not have the capacity to corral PI(4,5)P_2_, or other lipids, over the ∼0.1 s and ∼0.1 µm scales that we resolve – scales that are highly relevant to the assembly and regulation of these complexes. Rapid PI(4,5)P_2_ diffusion in the vicinity of these complexes will quickly dissipate any local enrichment of unbound lipid, even if it is locally synthesized. It therefore seems very unlikely that local PI(4,5)P_2_ enrichment serves as a platform to induce assembly of components such as clathrin coated structures or ER-PM contact sites *de novo*.

That is not to say that once these complexes begin to assemble, engagement of their effector proteins with PI(4,5)P_2_ will not enrich the lipid, and be crucial for growth and regulation of these complexes. Such effector-bound PI(4,5)P_2_ is invisible to our lipid biosensors. However, it is implicit from our data that local enrichment of PI(4,5)P_2_ must be driven by effector proteins, and not the other way around.

## Materials and Methods

### Cell Culture and transfection

HeLa (ATCC CCL-2) cells were grown in DMEM (low glucose; Life Technologies 10567022) supplemented with 10% heat-inactivated fetal bovine serum (Life Technologies 10438-034), 100 units/ml penicillin, 100 µg/ml streptomycin (Life Technologies 15140122) and 1:1000 chemically-defined lipid supplement (Life Technologies 11905031) at 37°C in a humidified atmosphere with 5% CO2.

For transfection, cells were seeded in 35 mm tissue culture dishes with 20 mm Number 1.5 cover glass apertures (CellVis) previously coated with 5 µg fibronectin (Life Technologies 33016-015). After 1 to 24 hours post seeding, cells were transfected with 0.5 µg of plasmid DNA coding for lipid biosensors. For overexpressed domains 0.5 µg of plasmid DNA were used.

Plasmids were pre-complexed with 3 µg lipofectamine 2000 (Life Technologies 11668019) in 200 µl Opti-MEM (Life Technologies 51985091) according to the manufacturer’s instructions. Cells were imaged 4-18 h post-transfection. For latrunculin-B treatment (AbCam 144291), cells with Septin2 endogenously labeled with sfGFP were incubated with 1µM Latrunculin-B for 30 min before imaging.

### Generation of endogenous tagged cell lines

HeLa cells endogenously tagged with split GFP were produced as described (Leonetti et. al, 2016) using a protocol we have described (Zewe et al., 2018). HeLa cells stably expressing sfGFP-1-10 were electroporated with single-stranded Homologous-directed repair (HDR) template (IDT) and precomplexed gRNA and Platinum Cas9 (Thermo Fisher). Sequences are provided in **table 5**. All HDR templates contained 70 bp homology-arms, the GFP-11 sequence, and a flexible linker in frame with the gene to be labeled (CGTGACCACATGGTCCTTCATGAGTATGTAAATGCTGCTGGGATTACAGGTGGCGGC). 48 h after electroporation, cells were sorted by FACS for GFP-positive cells.

**Table 5.**
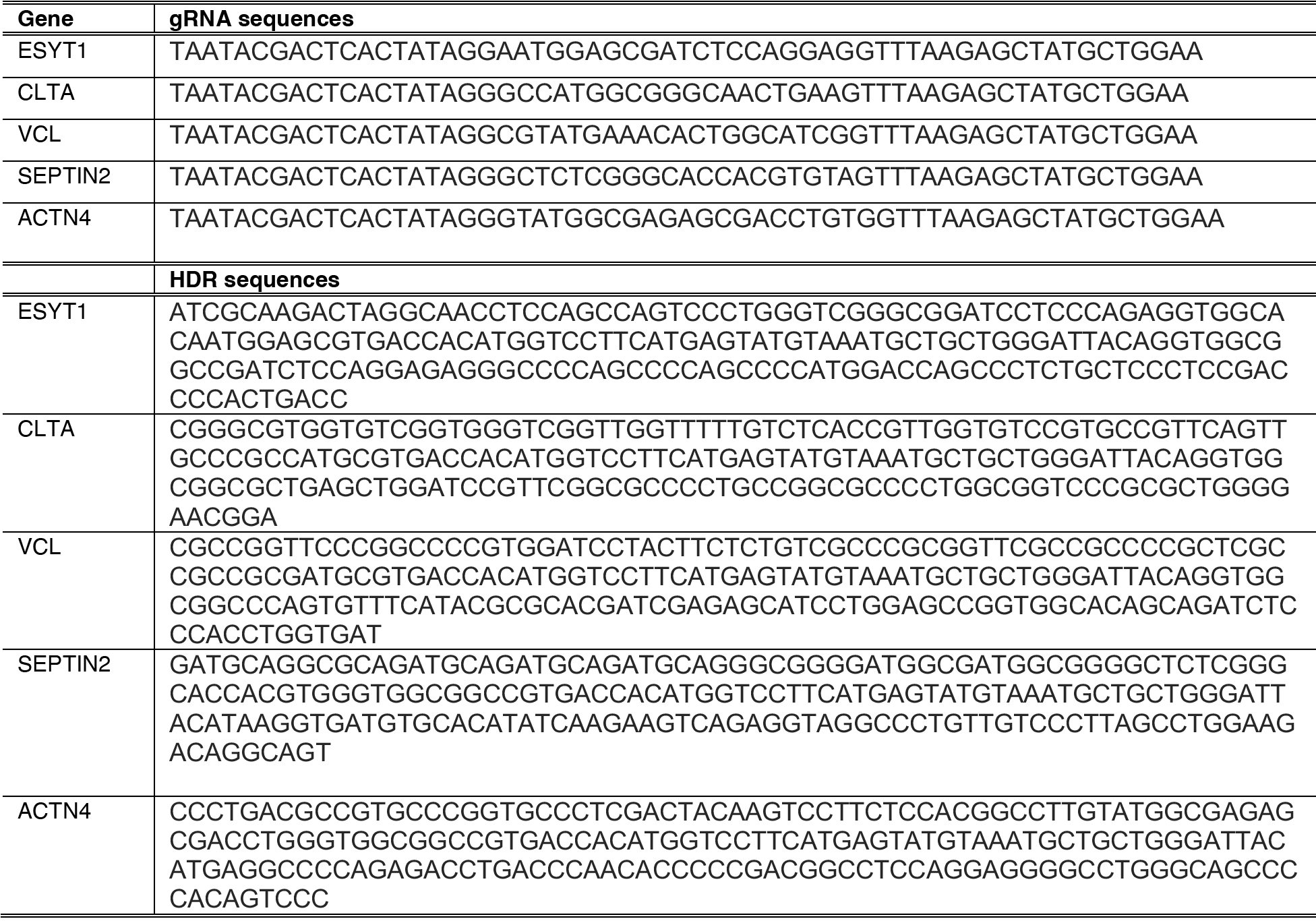
HDR and gRNA sequences for targeted genes

### Plasmids

pPAmCherry1-C1 (Addgene plasmid 31929) was a kind gift of Vladislav Verkhusha (Albert Einstein College of Medicine, New York, NY). The EGFP (*Aequorea Victoria* GFP presenting F64L and S65T mutations) was used to label plasma membrane domains. ARPC4 cDNA was obtained from GeneCopoeia^TM^. HIV-1-GAG fused with EGFP was purchased from Addgene (plasmid 80605). Ezrin plasmid was a generous gift of Adam Kwiatkowski (University of Pittsburgh School of Medicine, Pittsburgh, PA). All plasmids were verified by dideoxy sequencing. Plasmids were constructed using NEB HiFi assembly or standard restriction cloning. Sources and backbones are indicated in **table 6**.

**Table 6.** Plasmids used in this study

| Plasmid | Backbone | Insert | Reference |
| --- | --- | --- | --- |
| PAmCherry1-N1-Tubbyc | pPAmCherry-N1 | Mus musculus Tub (243-505): PAmCherry1 | This study |
| PAmCherry-N1-FRB-Tubbyc | pPAmCherry-N1 | Mus musculus Tub (243-505):MTOR(2021-2113):PAmCherry1 | This study |
| mCherry-N1-Tubbyc | pEGFP-N1 | Mus musculus Tub (243-505): mCherry | Quinn et al., 2008 |
| PAmCherry1-N1-PHPLCd1 | pPAmCherry-N1 | PLCD1v2(1-170):PAmCherry1 | This study |
| NES-PAmCherry-cPHx3 | pPAmCherry-C1 | PAmCherry1:PLEKHA1(169-329):GGSGGSGG: PLEKHA1(169-329): GGSGGSGG: PLEKHA1(169-329): | Goulden et. al., 2019 |
| PAmCherry1-C1-P4C | pPAmCherry-C1 | PAmCherry1: *L. pneumophila* SidC (608-773) | This study |
| PAmCherry1-C1-P4Mx2 | pPAmCherry-C1 | PAmCherry1: *L. pneumophila* SidM(546-647):SidM(546-647) | This study |
| PAmCherry1-C1-LACT-C2 | pmEos2-C1 | PAmCherry1: *Bos taurus* MFGE8 (274-431) | This study |
| PAmCherry1-N1-Lyn11 | pmEos2-N1 | LYN(1-11):PAmCherry1 | This study |
| ARPC4 | pReceiver-M98 | EGFP:ARPC4 | GeneCopoeia™ |
| EGFP-N-Ezrin | pmCherry-N1 | Mus musculus EZR(1-586):EGFP | This study |
| EGFP-C3-β-spectrin | pEGFP-C3 | EGFP:SPTB(1-2364) | This study |
| EGFP-N1-Lifeact | pEGFP-N1 | LifeAct:EGFP | Tamas Balla |
| EGFP-C1-FKBP-CYB5<sup>tail</sup> | EGFP-C1 | EGFP:FKBP:[GGSA]<sub>4</sub>GG:CYB5A(100-134) | This study |

### Microscopy

Cells were imaged in 2 ml FluoroBrite DMEM (Life Technologies A1896702) supplemented with 25 mM Hepes (pH7.4), 1:1000 chemically defined lipid supplement with 10% heat-inactivated fetal bovine serum. For fixed cell preparations, transfected cells were washed with warmed Phosphate-Buffered Saline (PBS) and immediately fixed with 0.2% Glutaraldehyde dissolved in PBS. After 15 min incubation, cells were rinsed 3 times with freshly diluted 10 mg/ml Sodium Borohydride dissolved in PBS. Finally, cells were washed twice with PBS and imaged.

All experiments were performed on a Nikon TiE microscope equipped with a TIRF illuminator, a 100X 1.45 NA plan-apochromatic oil-immersion objective and an Oxxius L4C combiner equipped with 405, 488, 561 and 640 nm lasers. Single molecule imaging was registered on a Zyla 5.5 sCMOS camera (Andor) with no pixel binning in a rolling shutter mode. Detection of single molecule and domains were recorded with a frame delay of 0.025 ms by using a sequential acquisition between green (488 nm laser line excitation for the EGFP tagged PM domains) and red (561 nm excitation for PAmCherry) on triggering mode controlled by NIS-Elements software. Final exposure for single molecule imaging resulted in 55 ms by using 30% laser power with 561 nm and from 0.5 to 3 seconds of 0.8% of 405 nm laser for photoactivation immediately before starting the experiment. Green channel was excited with 3 to 8% of 488 nm laser. Time lapse images were recorded in a 16 x 16 µm region of the PM for 30 seconds.

### Single Molecule Analysis using Thunderstorm

Spot size and brightness (**Fig. 1C and 1D**) was estimated with Fiji thunderstorm plugin (Ovesný et al., 2014). Fixed cells expressing PAmCherry1-N1-Tubby*c* were imaged using the same microscope settings as for live cell single molecule time lapses. Then, raw images were run in the thunderstorm plugin. Settings for localization of molecules were determined by using a wavelet filter with a local maximum method and an integrated Gaussian Point spread function.

For fluorescence intensity per spot, histograms of photon counts were generated with a bin size of 5 photons. For size of spots, histograms of 2 standard deviations of detection (sigma) were produced with a bin size of 4 nm.

### Analysis of the trajectories

Single molecule trajectories were generated using the open-source Fiji TrackMate (Tinevez et al., 2017). For single molecule detection a difference of Gaussians filter was used with an estimated diameter of 0.5 µm (i.e., an ∼8x8 pixel neighborhood). For trajectory assembly an implementation of a simple linear assignment problem (LAP) algorithm with 0.7 µm maximal distance for both linking and gap-closing was used. Coordinates of trajectories were exported as CSV files and analyzed using custom written code in Python to calculate trajectory displacements. For all analyses, only those trajectories longer or equal to 0.88 s (16 frames) were used.

Individual trajectories were analyzed through the mean-square displacement (MSD) (Vrljic et al., 2007). The trajectories were cut at a duration of 0.88 s and the MSD was calculated from the first 15-time lags. The diffusion coefficient was extracted from a power of law function using the 16-time lags of MSD plots:

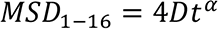

All independent displacements (r) from all trajectories were pooled to construct cumulative distribution functions at different time lags (iΔt) (Vrljic et al., 2002). The cumulative distribution function plots were constructed for the first 4-time lags. Unless otherwise mentioned, each curve was fit with equation 2, which considers two populations of diffusion coefficients:

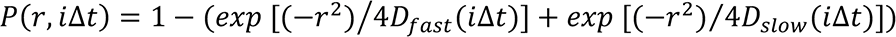

Where Dfast and Dslow is the diffusion coefficient for the fast and the slow population, respectively. Dfast > Dslow at all time lags. To analyze whether diffusion is Brownian or anomalous, the change of diffusion coefficient over time was evaluated for each population:

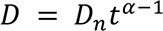

Where the exponent alpha denotes the grade of anomalous motion. Dn is either, Dfast or Dslow.

For Brownian motion the exponent alpha is close to 1. For subdiffusive and superdiffusive motion, alpha is less than 1 or greater than 1, respectively. Fitting was performed using Graphpad Prism 9.

### Analysis of diffusion in-domains

The trajectories were obtained through sequential acquisition (delayed time of ∼27 msec) between PM-domains (Green channel) and single molecule detection (Red channel). The expression of EGFP labeled domains was used to generate binary masks through à trous wavelet decomposition (Olivo-Marin, 2002) as described previously (Hammond et al., 2014).

To quantify diffusion coefficients inside domains, we used a moving window approach. A moving window was defined as a unit of analysis of a partial segment of trajectory with a length of 5 consecutive localizations (iΔt=5). The moving window is scanned through the trajectories step by step until the whole trajectory is covered. All those partial trajectories produced by the moving windows were categorized as an in-domain measurement if at least one localization fell inside of a domain. In the same way, all moving windows in which all localizations were outside of domains correspond to out-of-domain measurement. Each partial segment was used to construct cumulative distribution functions of radial displacements (**equation 2**). Diffusion coefficients for fast and slow populations were calculated using the first 4-time lags (**equation 3**). Additionally, all partial trajectories produced by the moving windows were analyzed by MSD (**equation 1**), and the median was graphed per cell.

Analysis of septin rings was performed exclusively in those domains that under the binary mask were completely closed rings. From trajectories, the number of crosses was counted. A crossing was defined as a segment produced by the Euclidean vector between localizations that move through the inner side of the ring or vice versa. For control, the crosses were counted in randomized binary masks produced by flipping vertically, horizontally, and vertical-horizontally the same masks with septin rings produced from experimental data.

### Turning angles

The turning angles were quantified from the resultant angle between 2 consecutive vectors along the trajectories. Only trajectories longer than 16 frames were used. To measure the turning angles in domains, only whole trajectories moving through domains (same binary masks from previous section) were considered for the analysis. Here, an in-domain measurement was established if at least one localization in a vector is positioned in the coordinates corresponding to a domain. An out-of-domain measurement was all the turning angles from segments of vectors moving out of domains.

### Statistical Analysis

All statistical analyses were performed using Graphpad Prism 9. Trajectories were analyzed as above, and the mean values were computed based on the total trajectories recorded from individual cells. Note that data was collected from ≥ 3 independent experiments, though variability among cells in each experiment was greater amongst cells in a given experiment than it was between experiments. Thus, we define the cell as the unit of biological variability. In the data collected from individual domains (reported in **figures 4-8**), variability in the number of trajectories interacting with domains led to a small subset of cells with highly divergent estimates of diffusion coefficients and alpha for the slow population, which skewed the mean. Therefore, we subjected all these data to outlier analysis using the ROUT method (Matulsky and Brown, 2006), setting a maximum false discovery rate (Q) of 0.1%. Any cell with a detected outlier in any parameter was excluded from further analysis. Statistical tests were performed as described in the figure legends for each experiment and detailed in **tables 1-4**.

## Contributions

JP, ACC and GRVH performed single molecule tracking experiments and analyzed data. JP, JPZ and RCW prepared gene edited cell lines. GRVH acquired funding and wrote the manuscript. All authors edited the manuscript.

## Acknowledgements

The authors are grateful to Scott Hansen (University of Oregon, Eugene OR) for critical reading of the manuscript and helpful suggestions.

